## Supplementary text and figures 1-15 for "Comparative whole-genome approach to identify bacterial traits for microbial interactions"

**This PDF file includes:**

Supplementary text

Supplementary figures 1 to 15

**Other supplementary materials for this manuscript include the following:**

Supplementary tables 1 to 10

Supplementary files 1 to 5

### An overview of approaches for functional genome classification

Over the last >20 years, since genome sequencing became widespread, many studies have aimed to classify organisms based on the functions encoded in their genomes (see Supplementary Table 1 for a detailed yet likely not comprehensive list). Below, we briefly summarize these studies, and highlight where the approach we utilize here builds upon these studies and provides new insights.

The nineteen studies detailed in Supplementary Table 1 can be divided along two main aspects: the type of genomic information analysed (genomes VS metagenomes) and the resolution of the functional annotations considered (single genes VS traits or functional categories). Genome-based studies (including both draft and complete genomes) mainly focused on specific taxa (e.g. *Bacillus*, *Clostridia*, *Roseobacter*) [1–5], although two notable exceptions focused on a wide diversity of marine bacteria [6,7]. Based on their genomes, marine bacteria can be classified into two main groups – oligotrophs, which are often highly abundant, and copiotrophs, which are often less common but can grow rapidly in energy-rich environments [6]. These two groups differ in the size of their genomes (which are much smaller and more streamlined for the oligotrophs) and the relative abundance of specific broad-scale functions (e.g. periplasmic, outer-membrane or extracellular proteins), functional categories (e.g. COG categories such as motility or signal transduction) or specific genes groups (COGs such as COG0583 – transcriptional regulator) [7]. More detailed studies of specific taxa (e.g. Roseobacters) often highlighted relatively large functional differences within specific clades, which often were not congruent with phylogeny [5]. Notably, metagenome-based studies, or those analysing genomes from single cells, often encompassed a wider taxonomic diversity [8–11]. Such approaches allowed to describe an unprecedented functional uniqueness of bacterial and archaeal single-cell amplified genomes (SAGs) in tropical and subtropical ocean, which bore numerous pathways involved in light harvesting and secondary metabolite biosynthesis [11]. Similarly, the analysis of metagenome-assembled genomes (MAGs) highlighted that certain COGs involved in saccharide and lipids biosynthesis, nitrate and sulfate reduction, as well as $CO_2$ fixation were specifically enriched in marine prokaryotes inhabiting polar regions [10]. However, due to the often incomplete nature of MAGs and SAGs, such studies also have a lower functional resolution (e.g. missing less common function/genes), and do not take into account the absence of specific traits (e.g.. in [10,11]).

As noted above, functional annotation can be performed at multiple levels of resolution, from very broad-scale functions (e.g. “extracellular proteins”) to individual genes. Overall, the majority of the studies presented in Supplementary Table 1 focused on gene-level annotation [1–4,8–13]. Analysing genomes or metagenomes at the single-gene level enabled the resolution of fine differences in the functional capacity between bacteria, e.g. defining ecotypes [2] or revealing limited clonality in bacterial communities [11], but often at the cost of a clear overview of the processes and/or pathways actually encoded. In contrast, studies that characterized genomic information in more coarse functional categories (e.g. COGs or COG categories) often highlighted relevant features such as cell motility, sensory systems or secondary metabolite production that characterized bacterial lifestyles [7] or environmental preferences [3,5,6,10,14]. A trait-based analysis was developed to characterize the capacity of different bacteria in terms of multiple substrates utilization, oxygen requirement, morphology, antibiotic susceptibility, or proteolysis. However, the workflow was based on a commercial platform (GIDEON) and mainly focused on medical-related phenotypes and bacteria (belonging to Gammaproteobacteria, Firmicutes, Bacteroidetes, Actinobacteria) [15].

In our study, we chose an approach that builds upon previous knowledge but differs in two main ways. Firstly, our analysis encompassed a wide taxonomic diversity of marine bacteria (421 strains, 213 genera), using only complete genomes to minimize false negative occurrence of genetic traits. Secondly, we chose an intermediate functional resolution to annotate these genomes – that of genetic traits, defined here as the presence of complete gene pathways (e.g. KEGG modules, pathways for biosynthesis of secondary metabolites and phytohormones, vitamin and siderophore transporter). This resolution is more detailed than that of COG functions or specific COGs, providing a direct link between gene annotation and cell metabolism of specific compounds, while covering a wider range of genetic traits with a specific focus on bacteria interaction with other microorganisms. By defining genetic traits and linking them into Linked Trait Clusters (LTCs), and by using such traits to cluster genomes into Genome Functional Clusters (GFCs), this framework offers an efficient way for translating genomic information into physiologically- and ecologically-relevant traits, and for classifying bacteria into groups which we propose perform similar functions.

#### Remarks on the annotation pipeline

A relevant aspect of our analysis which needs to be kept in mind: we included only closed bacterial genomes (i.e. a single, high quality sequence of each DNA molecule such as chromosome or plasmid) or high-quality draft genomes (estimated by using CheckM, see method section). The rational was to provide a comprehensive description of the full functional potential of pelagic marine bacteria which requires high-quality genomes to achieve the best information possible [16]. Gene annotation is *per-se* a challenging step, in particular when it deals with environmental genomes for which many genes are still unknown and, therefore, cannot be properly annotated (in our analysis ~63% of the predicted coding sequences were annotated).

A further step to improve cross-comparability among genomes was to re-annotate all of them using a standardized pipeline. We developed a trait-based workflow which, instead of looking at the level of single annotated genes, detects the presence of complete genetic traits aiming to a more robust prediction of the inferred metabolic potential. The majority of the annotated traits were KEGG modules (KMs; ~87% of total traits, Figure 1). KMs represent defined functional units (e.g. the glycolysis pathway; Supplementary Fig. 1b) and their completeness was assessed taking into account potential annotation issues (Supplementary Fig. 1C; more details in the benchmarking section below). The genome functional profiles were further enriched with the annotation of other genetic traits using specific tools, e.g. secondary metabolites (antiSMASH), transporters (BioV suite), phytohormones production (KEGG pathway map01070), vibrioferrin production and tranport, as well as the degradation of dimethylsulfoniopropionate (DMSP), 2,3-dihydroxypropane-1-sulfonate (DHPS) and taurine (manual annotation; see Material and Methods; Supplementary Fig. 1d).

Additionally, the presence of a complete genetic trait did not necessarily translate into an expressed phenotype. The correlation between gene content and phenotype has been shown for some traits (e.g.

motility [17]), however, several genetic traits may be not constitutively active. Their expression could be under fine regulatory controls and the relevant phenotypes would manifest only under specific environmental and/or physiological conditions.

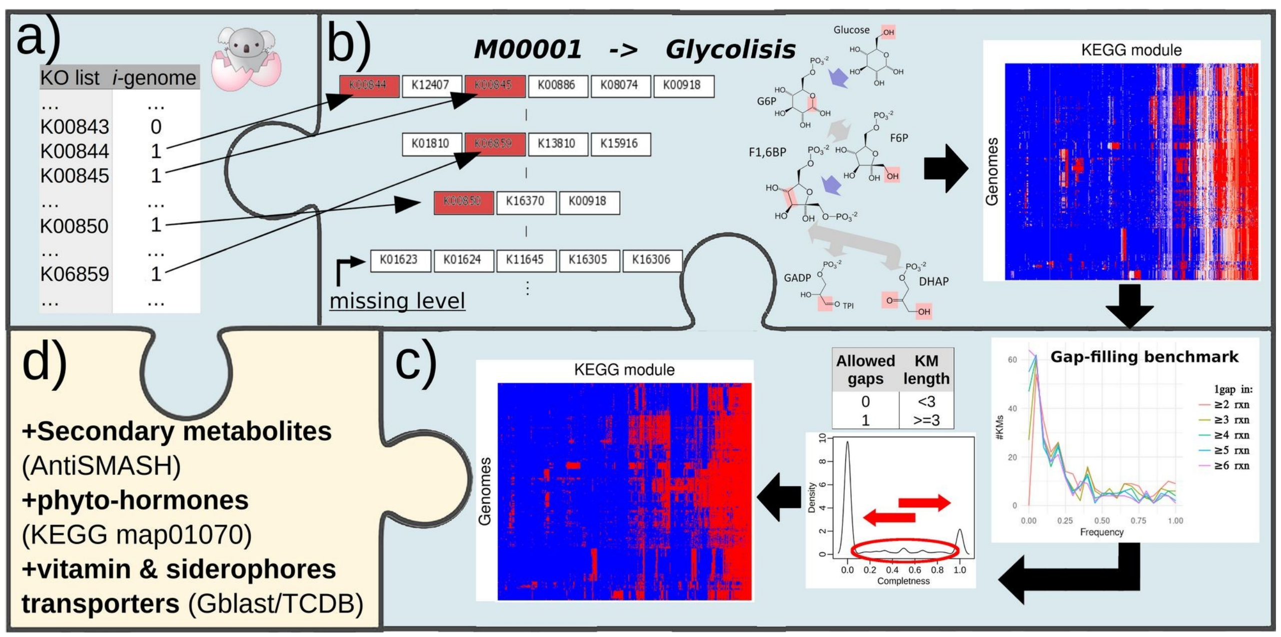


Supplementary Fig. 1: Annotation workflow for the identification of genetic traits in genomes (a-c). Annotated KEGG orthologies (a) were recombined in all known KEGG modules (KM; b) and labelled as present or absent using a custom R script. The script, taking into account some completion rules, generates a presence/absence matrix (c). Further annotations were performed using antiSMASH for detection of secondary metabolites, KEGG orthologues of the pathways map01070 for detection of phytohormones and Gblast against the Transporter Classification Database (TCDB) to identify B vitamins and siderophore transporters (d).

### GFCs and their taxonomy

The genome clustering analysis retrieved a total of 47 genome functional clusters (GFCs). As shown in Supplementary Fig. 3a, most of these GFCs included only genomes of the same phylum (40), and fewer than 3 different families (10 with 1 and 18 with 2). At the genus levels, more than half of the GFCs included 3 or more genera. From the opposite perspective, at the taxa level, ~35% of the phyla were represented by 2 or more GFCs, while nearly all genera (~94%) were represented by a single GFC (Supplementary Fig. 3b).

We found that some GFCs represented group of organisms with a defined ecology and life history. For example, GFC 2 comprised all genomes of the order SAR11 (Pelagibacterales) (Supplementary Table 3), defining a group of highly abundant taxa with streamlined genomes adapted to thrive under oligotrophic conditions [18,19]. The GFCs 15 and 36 were to a large extent consistent with previous ecological and genomic studies on Cyanobacteria, with GFC 15 comprising *Synechococcus* and low-light type IV *Prochlorococcus* strains, while GFC 36 grouped exclusively *Prochlorococcus* strains of high-light type I-II and low-light I-III (reviewed by [20]). Genomes belonging to the family Vibrionaceae were clustered in two different GFCs (25 and 47). GFC 25 grouped known host of zooplankton (e.g. *Vibrio alginolyticus;* [21]), as well as other non-pathogenic strains (e.g. *V. furnissii* and *V. natriegens;* [22,23]). GFC 47 included several pathogenic strains of more generalist *Vibrio* species characterized by a wide range of aquatic hosts (e.g. *V. splendidus*; [24]), as well as a few human pathogens (*V. cholerae* and *V. vulnificus*; [22]). Along with *Vibrio* genomes, GFC 47 contained also genomes from additional taxa (e.g., *Photobacterium* (3 strains) and *Psychromonas* (2 strains)) which are also potential pathogens or gut endobionts of crustacean and marine snails [25,26].

We note, however, that the GFC analysis did not reproduce some aspects of high-resolution functional differentiation between closely related bacteria, e.g. between specific high-light ecotypes in Prochlorococcus (which share the “high light” surface niche but vary in their temperature or nutrient optima) [20] or between different species of Alteromonas, that are also supposed to inhabit slightly different niches [27].

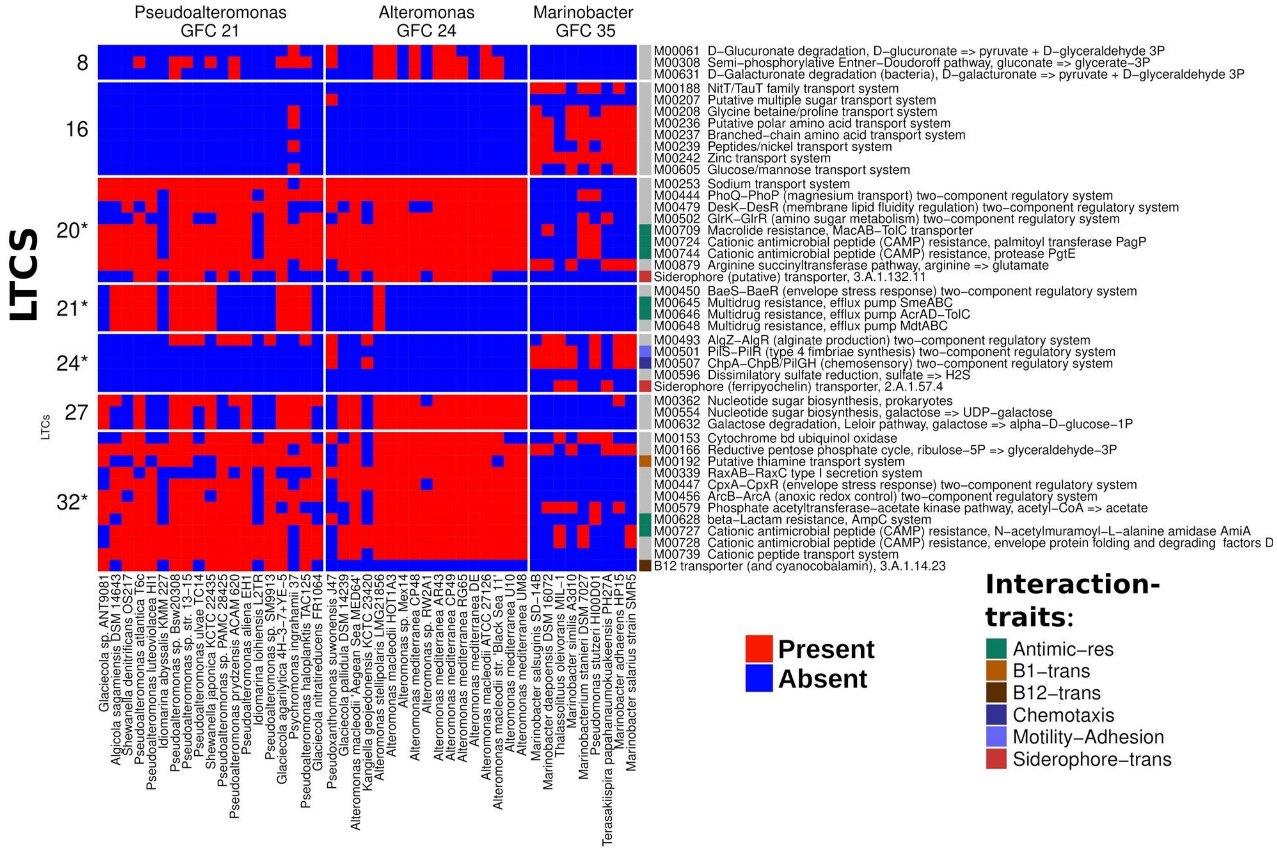


Supplementary Fig. 2: Insight analysis on the different genetic traits possessed by *Pseudoalteromonas*, *Alteromonas* and *Marinobacter*. The columns represent the different genomes, and the rows different genetic traits, combined into Linked Trait Clusters (LTCs, as presented later in the main text). The asterisk on the LTC names indicates that a given LTC contains also one or more interaction traits. All three GFCs share the the LTCs 2, 3*, 4, 5*, 7*, 9*, 10*, 11* and 13*; please refer to Supplementary Table 4 for more details about these LTCs.

Similarly to [28], the taxonomic coherence of each GFC was calculated based on the "local" taxonomic coherence score ($TC_{GFC}$; Supplementary Fig. 3c):

$$TC_{GFC} = N_{GFC}/N_{taxon}$$

Where $N_{GFC}$ is the number of genomes grouped in a GFC and $N_{taxon}$ is the total number of genomes (included in our genome atlas) belonging to the last common ancestor of the GFC. The level of taxonomic coherence of a GFC corresponds to the taxonomic rank of the last common ancestor. A $TC_{GFC}$ of 1 indicates that all genomes belonging to the last common ancestor are included in the respective GFC which is considered to be monophyletic. A $TC_{GFC} < 1$ indicates instead a paraphyletic GFC. As the current genome availability doesn't allow for a uniform coverage of all bacterial taxa, a possible issue leading to taxonomic promiscuity of the paraphyletic GFCs might be related to the inclusion of "singleton" taxon (i.e. a single genome that represents a different taxon). To avoid interpretation biases due to these singletons, we excluded a maximum of one singleton genome per GFC in the computation of the taxonomic coherence (see Supplementary Fig. 3d). Nevertheless, the paraphyletic GFCs that included most genomes also had more evenly represented taxa, like GFCs 6, 10, 38 and 40 that grouped different taxa with > 2 genomes each.

Polyphyletic GFCs included organisms from multiple phyla, however these GFCs group genomes of organisms were isolated from extreme environments (e.g. thermal vents or hyper saline environments; GFCs 33 and 41) and marine sediment (GFC 17). These genomes were added in the analysis as outer groups and their taxonomic and functional diversity was not adequately covered. Nonetheless, extreme environments are hotspots for gene exchange processes e.g. horizontal gene transfer that, in turn, favours functional convergence even between distantly related organisms [29].

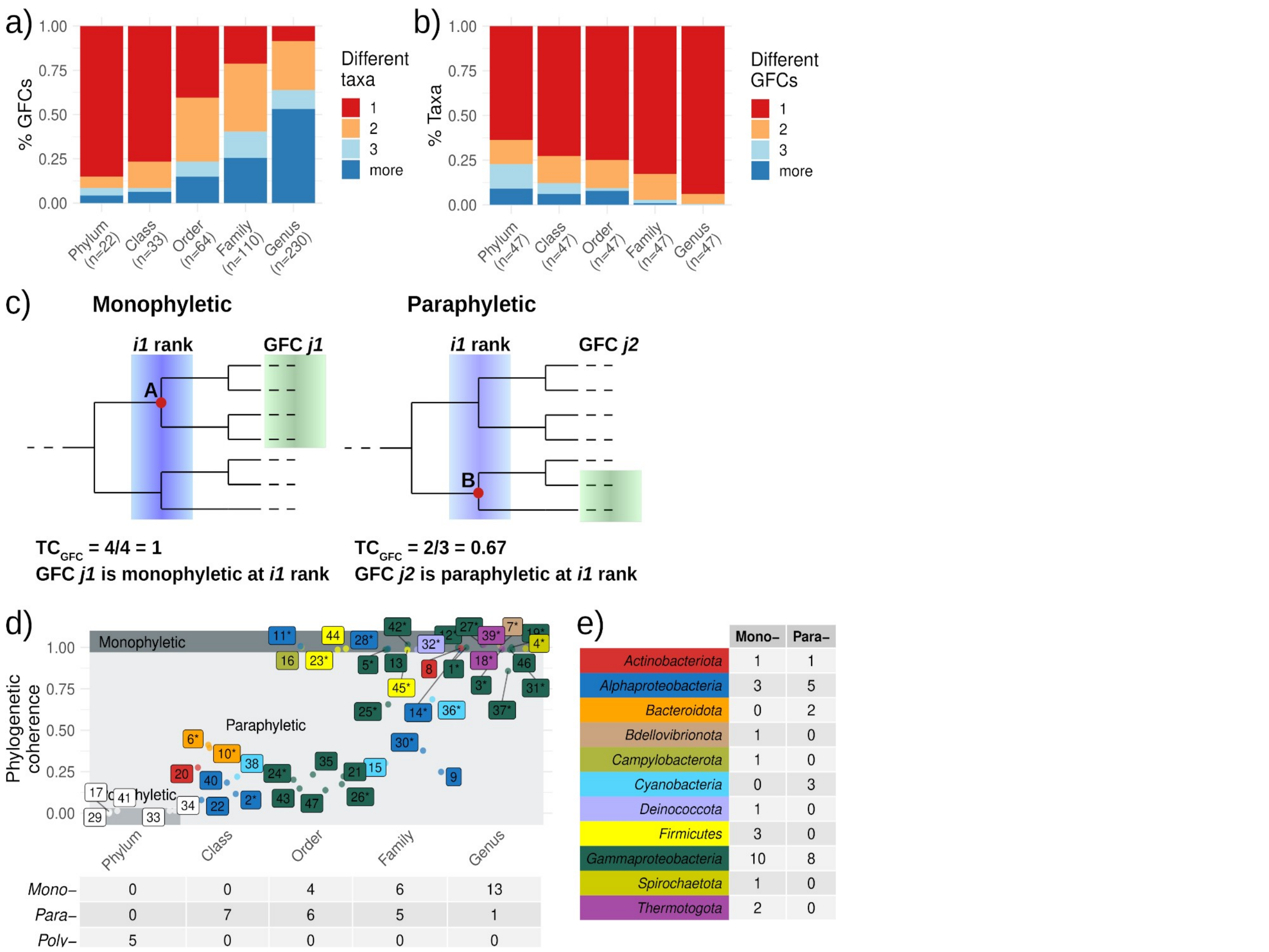


Supplementary Fig. 3: (a) Percentage of GFCs including 1, 2, 3 or more different taxa for each taxonomic rank. (b) Percentage of taxa included in 1, 2, 3 or more different GFCs for each taxonomic rank. (c) Schematic example of calculating the taxonomic coherence score in the genome functional clusters (GFCs) ‘j1’ and ‘j2’ carried out at the taxonomic rank ‘i1’. GFC ‘j1’ is monophyletic at that rank as it includes all genomes belonging to the related taxon ‘A’ (4/4), while GFC ‘j2’ is paraphyletic as it includes only some genomes of the related taxon ‘B’ (2/3). (d) Highest taxonomic coherence scores of each GFC separated by taxonomic ranks. Table underneath x-axis shows the amount of mono-, para- or polyphyletic GFCs for each rank; GFCs containing 1 “singleton” genome belonging to a different clade are marked with an asterisk; dots are colour coded according to their taxonomy as shown in panel (e), i.e. table summarizing the number of mono- and paraphyletic GFCs across different taxa.

### Mapping of genomes to a coastal and a pelagic time series

Genome mapping stringency was tested by using different thresholds of sequence identity. In the coastal time series, >90% of amplicon sequences mapped to unique GFCs (i.e. specificity = 1) with the 100% identity threshold, while in the pelagic time series that was already the case at the 97% identity threshold (Supplementary Figures 4a and 5a).

The average amount of mapped sequences in coastal communities varied between 22.9% and 18.3% at 100% and 97% sequence identity, while it was ~19% for the less stringent thresholds. The decrease in mapped sequences with lower mapping stringency was due to a higher share of promiscuous mapping (specificity <1) and such sequences were discarded from the analysis (Supplementary figures 4). The average number of mapped operational taxonomic units (OTUs) varied between 5.6% and 19.4% at 100% and 97% of sequence identity, but increased to ~30% for the less stringent thresholds. Among the mapped sequences and OTUs, the majority belonged to heterotrophic bacteria (13.1% - 17.2% on average, maximum 42.9%) such as Bacteroidota and Gammaproteobacteria, while only a smaller fraction mapped to Pelagibacterales (2.9% - 5.0% on average, maximum 17.6%) and Cyanobacteria (0.3% - 1.2% on average, maximum 7.3%).

In the pelagic time series, the average amount of mapped sequences varied between 13.9% and 34.6% at 100% and 97% sequence identity respectively, while it was ~45% for the less stringent thresholds. In this case, a lower mapping stringency did not affect the mapping specificity and increased the number of mapped sequences (Supplementary figures 5). The average number of mapped operational taxonomic units (OTUs) varied between 3.3% and 12.0% at 100% and 97% of sequence identity respectively, but increased to 30.6% at the 86.5% identity threshold. Among the mapped sequences and OTUs, the majority belonged to heterotrophic bacteria (7.8% - 24.2% on average, maximum 76.9%) such as Gammaproteobacteria, while only a smaller fraction mapped to Pelagibacterales (0.7% - 18.5% on average, maximum 45.4%) and Cyanobacteria (5.3% - 5.6% on average, maximum 19.4%). Moreover, as the samples were size fractionated, there was a clear distinction between the small filter pores (0.22 µm), with the majority of amplicon sequences mapping to GFCs which grouped Pelagibacterales and Cyanobacteria genomes (typical free-living taxa), and the large filter pores (11 µm), with the majority of sequences mapping GFCs which grouped other heterotrophic bacteria.

Using the temporal deconvolution analysis performed by Martin-Platero and colleagues for the coastal time series [30], we attempted a validation of the GFC concept, i.e. the genomes grouped in the same GFC have coherent functional profiles. Based on this definition, we bacteria belonging to the same GFC should display similar temporal trend in the environment as they are more likely to respond in the same way to environmental and biotic cues. We compared the frequency interaction scores of OTU pairs (or 16S phylotypes) mapped to a same GFC against the frequency interaction scores of OTU pairs mapped to different GFCs. The analysis showed that OTU pairs mapped to a same GFC had higher frequency interaction score, regardless of the identity threshold considered for the mapping (Supplementary Figure 6), indicating that such OTU pairs have synchronous temporal dynamics. Therefore, GFCs do actually partition bacterial diversity into groups with coherent functional potential and, likely, similar ecological niches.

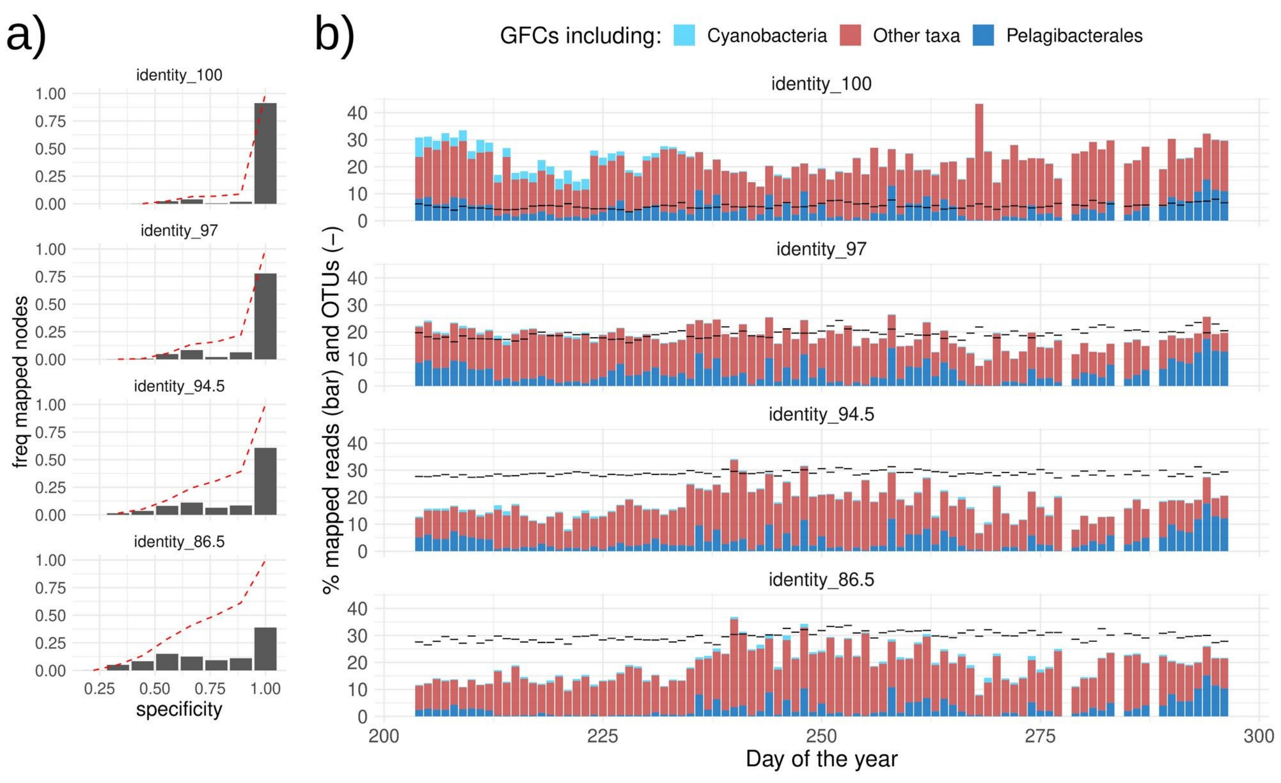


Supplementary figures 4: (a) histogram showing the frequency of specificity score computed across amplicon sequences of the coastal time series. Specificity score was calculated on the blast hits above the relevant identity threshold. Dashed red line shows the cumulative distribution of the frequency (b) Bar plots of the coastal site showing, at each identity threshold, the percentage of reads and OTUs that specifically (i.e. specificity = 1) mapped to any of the GFCs.

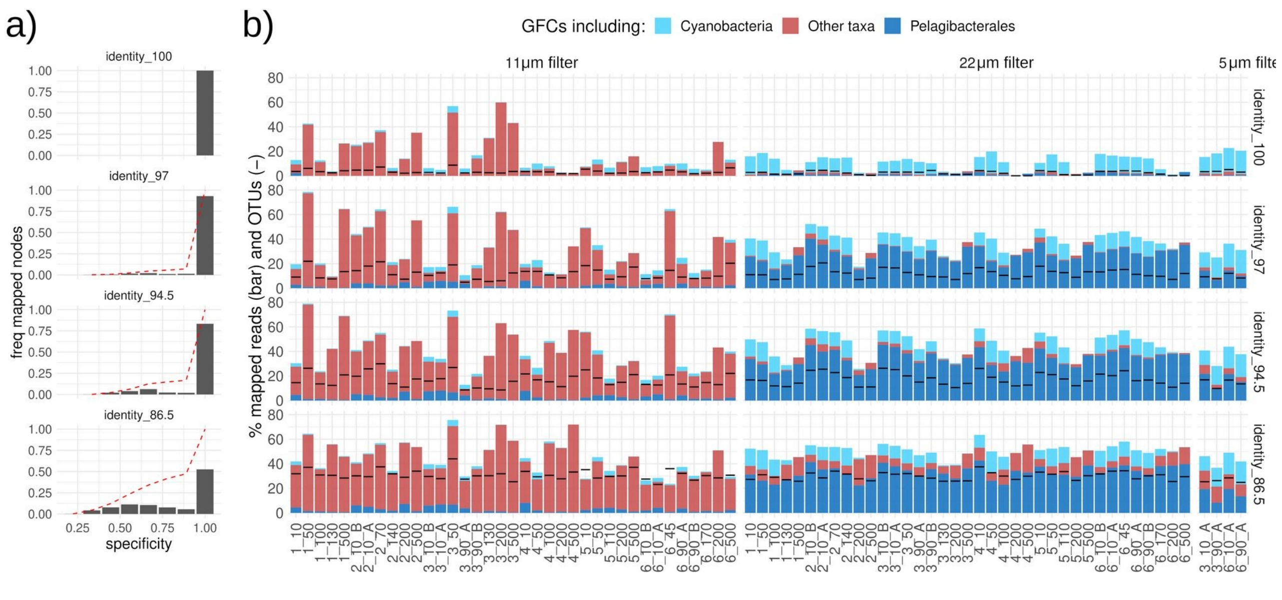


Supplementary figures 5: (a) histogram showing the frequency of specificity score computed across amplicon sequences of the pelagic time series. Specificity score was calculated on the blast hits above the relevant identity threshold. Dashed red line shows the cumulative distribution of the frequency (b) Bar plots of the pelagic site showing for each identity threshold, the percentage of reads and OTUs that specifically (i.e. specificity = 1) mapped to any of the GFCs. Sample names indicate the campaign and the depth at which samples were collected.

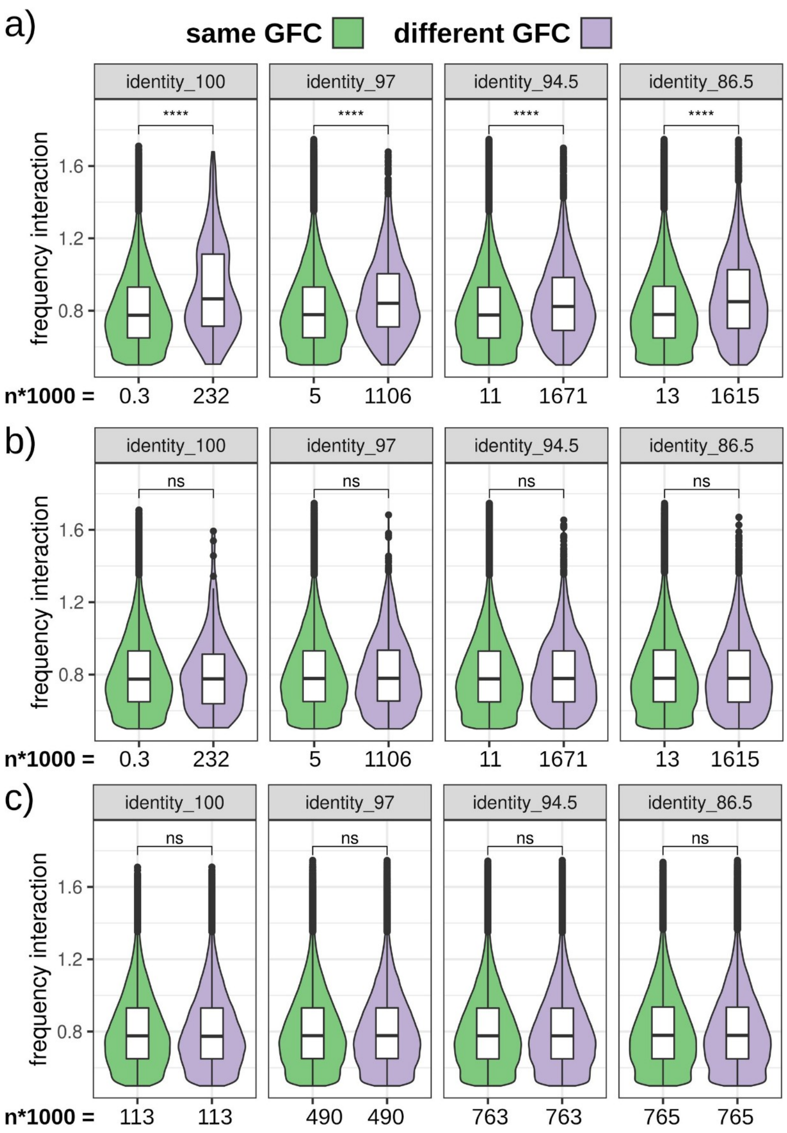


Supplementary figures 6: Violin plots showing the distribution of the frequency interaction score in OTU pairs mapped at different identity thresholds. (a) Comparison between OTU pairs in which both OTUs mapped to the same GFC or not. (b) Comparison between OTU pairs randomly assigned into two groups using the same group sizes of plot (a). (c) Comparison between OTU pairs randomly assigned into two groups of equal size.

### Genomes' enrichment in interaction traits

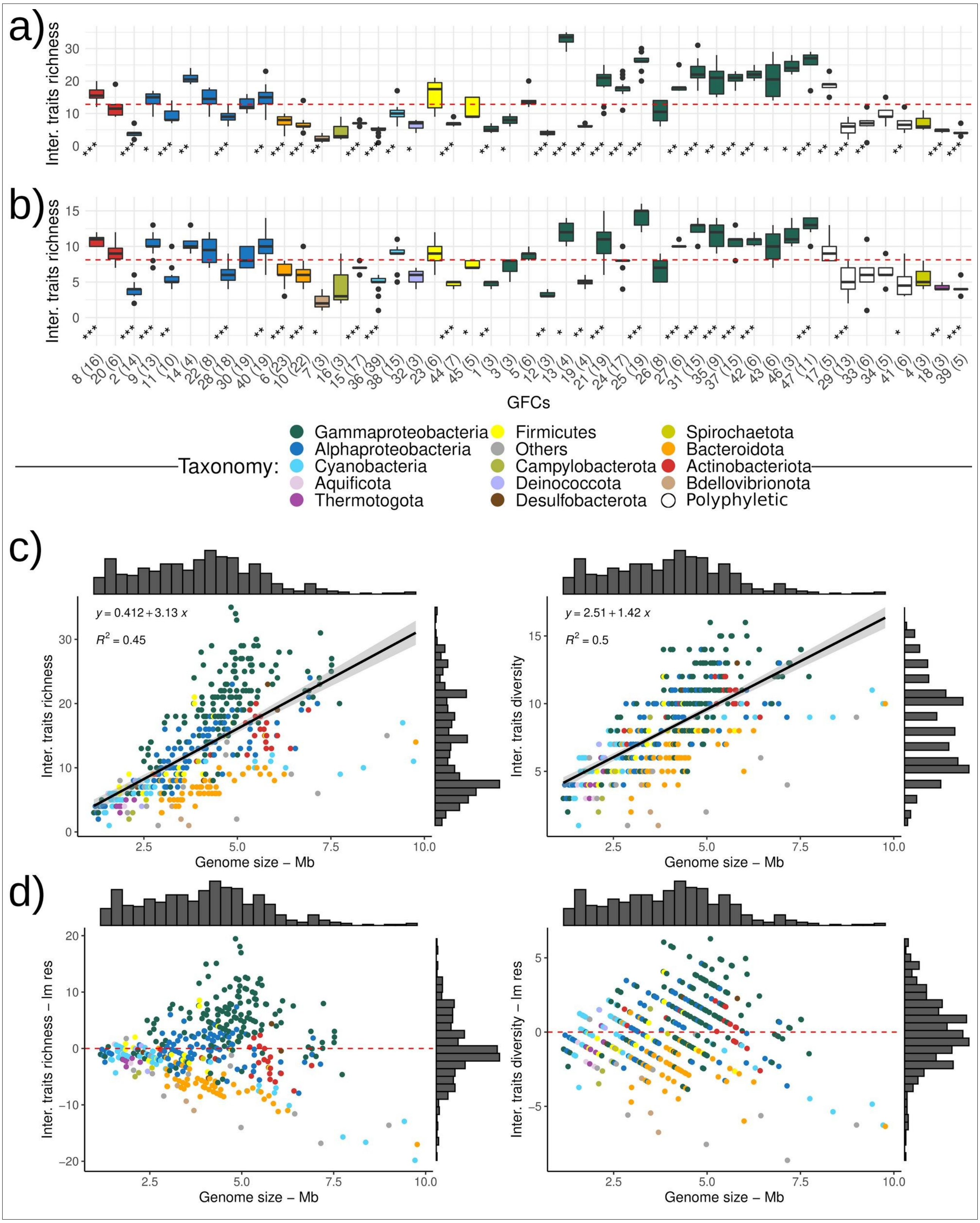
a)
b)
Inter. traits richness
GFCs
Taxonomy:
Gammaproteobacteria
Alphaproteobacteria
Cyanobacteria
Aquificota
Thermotogota
Firmicutes
Others
Campylobacterota
Deinococcota
Desulfobacterota
Spirochaetota
Bacteroidota
Actinobacteriota
Bdellovibrionota
Polyphyletic
c)
y = 0.412 + 3.13 x
R2 = 0.45
y = 2.51 + 1.42 x
R2 = 0.5
Inter. traits richness
Inter. traits diversity
Genome size – Mb
d)
Inter. traits richness – lm res
Inter. traits diversity – lm res
Genome size – Mb

Supplementary Fig. 7: For each Genome functional clusters (GFCs), the box-plots show the trait richness (i.e. total number; a) and the type richness (i.e. different types, visualized as different coloured squares in Fig. 2; b) of the interaction traits annotated in the grouped genomes. For each GFC, a t-test was performed to assess for a significant enrichment or depletion of interaction traits in comparison to the mean value of trait richness and diversity across all genomes (dashed red lines). (c) Linear regression models between genome size and interaction trait richness and diversity. (d) Plot of the residuals of the linear regression models (lm res). Genomes (i.e. dots) above the dashed red lines encode a higher number or a higher diversity of interaction traits than expected based on their genome size, while genomes underneath the lines bear less interaction traits than expected.

### Directionality of B vitamin transporters

Genomes with a flexible strategy could potentially act also as “source” for certain vitamins and represent key players in the vitamin market (e.g. [31–33]). However, the transport directionality can be reliably assigned only to specific transporter families (http://www.tcdb.org/superfamily.php) and we could identify only efflux-transporters for vitamin B1 (see Supplementary table 4 for directionality annotation) which were encoded in Alphaproteobacteria and Gammaproteobacteria genomes (Supplementary Fig. 8b). Moreover, it has to be kept in mind that B vitamins are water soluble molecules and could passively diffuse from a producing cell [34].

### Combinations of vitamin traits in specific GFCs

Biosynthetic pathways and related transporters have been identified also for other vitamins, e.g. biosynthetic pathways for vitamins $B_2$ (grouped in LTC 5, Supplementary table 7), $B_6$ (LTC 11), E (LTC 22) and $K_2$ (LTC 23), or the transporter for vitamin $B_3$ (LTC 29 and 30). Some of these vitamins are known to be exchanged during microbial interactions (e.g. [33,35]), however, the capabilities to produce and transport these vitamins were not consistently identified across genomes, and no clear pattern of bacterial strategy could be drawn for such vitamins (e.g. flexible/consumer/independent).

As shown in Fig. 2c, some of the most frequent combinations of synthesis and uptake of vitamins $B_1$, $B_{12}$, and $B_7$ appeared more often in certain taxa than others. These patterns suggested the existence of taxon-specific evolutionary strategies for handling these B vitamins. To test for this notion, we carried out an indicator species analysis as implemented in the function *multipatt* (func = "r.g", duleg = F, max.order = 5; package indicspecies) [36,37]. The function is designed to identify one or multiple species that can be used as indicators for certain habitats because of their strong species- habitat association. We therefore performed an analysis using all the B vitamins strategies (Fig. 2c and Supplementary Fig. 8a) instead of species, while GFCs were used instead of habitats. We didn't include GFC with < 3 genomes (i.e. only GFC 7 was excluded) and, as more than one GFC might share the same strategy, we allowed combinations of up to 5 different GFCs. Moreover, only associations with a p-value adjusted for false discovery rate < 0.05 were considered.

As shown in Supplementary Fig. 9, for 22 different B vitamin strategies (out of 51 possible ones) we identified a significant association with at least one of 43 different GFCs (out of 47). Except for Cyanobacteria, all other taxa had more than one associated strategy partitioned across different GFCs. Sometimes, different GFCs of the same taxon possessed the same strategy and in other cases the same strategy was present in GFCs of different taxa.

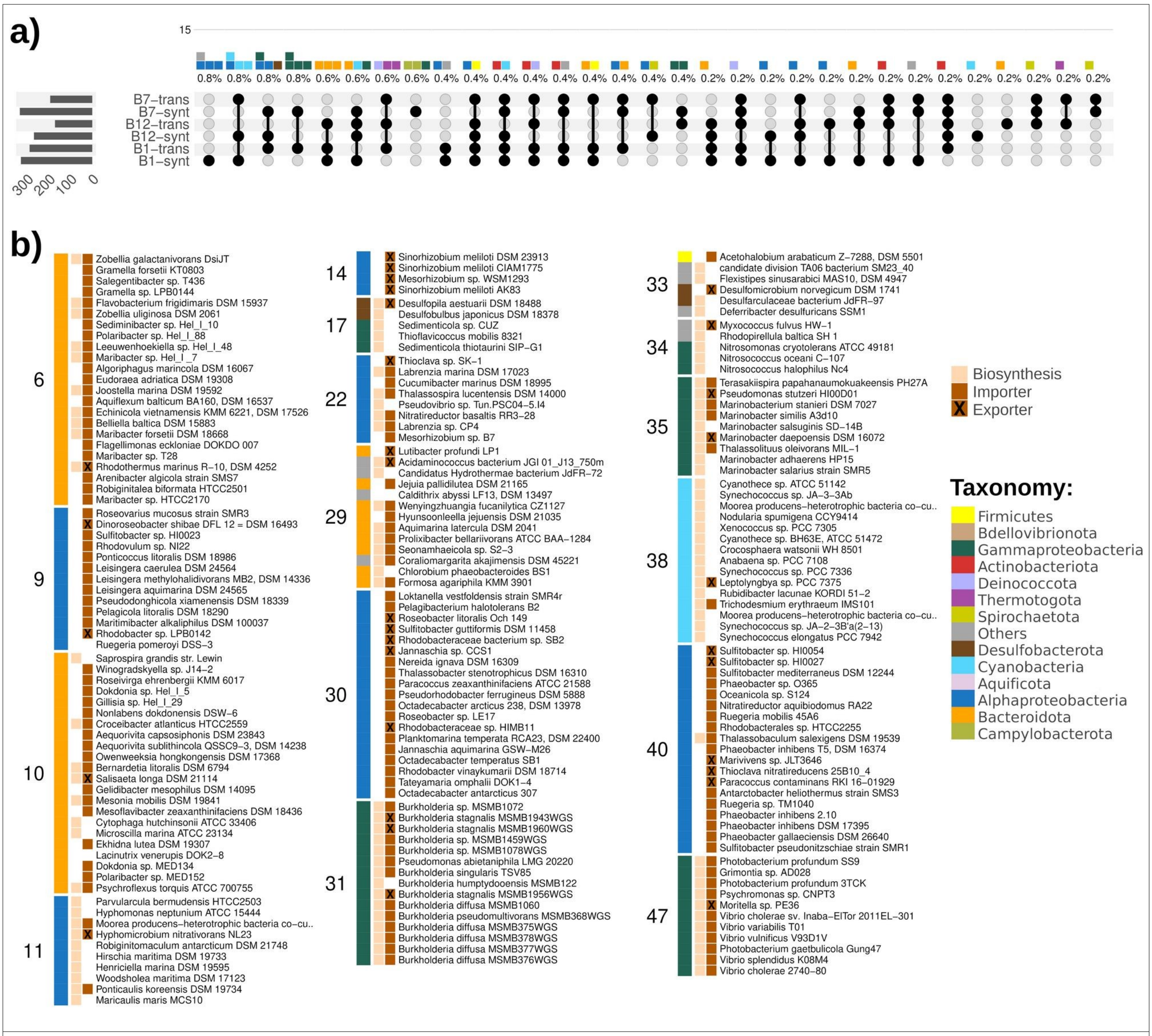


Supplementary Fig. 8: (a) Plot of intersecting sets showing the least abundant configurations of genetic traits related to production and/or transport of vitamins $B_1$, $B_{12}$, and $B_7$ (abundant configurations are shown in Fig. 3c) . The left bar chart indicates the total number of genomes for each trait, the dark connected dots indicate the different configurations of traits and the waffle bar chart indicates the number (and percentage) of genomes provided with such a configuration; each piece of a waffle bar represents a genome and it is coloured according to the taxon. (b) Insight into vitamin B1 flexibles genomes grouped in genome functional clusters (GFCs); GFCs colour bars correspond to the coherent taxonomy of the grouped genomes.

222

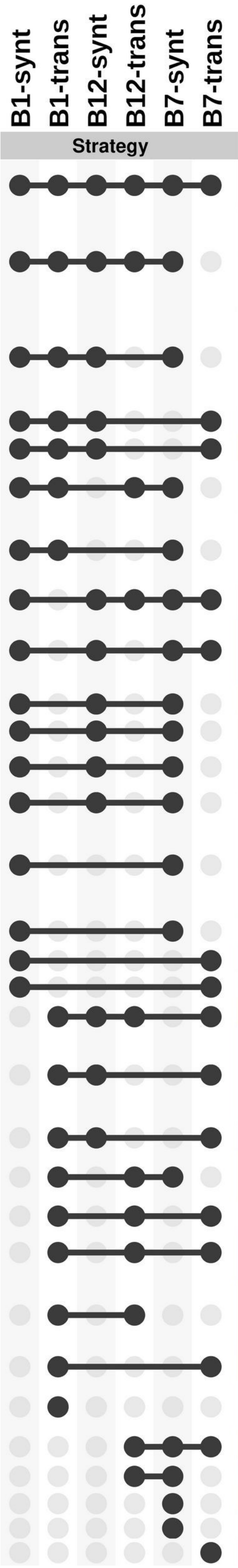


| Taxon | GFC | Coherent taxon | IndVal | p–adj |
|---|---|---|---|---|
| Actinobacteriota | 8 | g__Micromonospora | 0.47 | 0.011 |
| | 20 | c__Actinomycetia | 0.2 | 0.011 |
| Gammaproteobacteria | 37 | g__Shewanella | 0.48 | 0.0017 |
| | 25 | f__Vibrionaceae | 0.43 | 0.0017 |
| | 47 | o__Enterobacterales | 0.41 | 0.0017 |
| | 21 | o__Enterobacterales | 0.19 | 0.0017 |
| Gammaproteobacteria | 31 | g__Burkholderia | 0.6 | 0.0017 |
| | 35 | o__Pseudomonadales | 0.27 | 0.0017 |
| | 42 | f__Cellvibrionaceae | 0.19 | 0.0017 |
| | 43 | o__Pseudomonadales | 0.19 | 0.0017 |
| Alphaproteobacteria | 22 | c__Alphaproteobacteria | 0.48 | 0.016 |
| Firmicutes | 23 | o__Bacillales | 0.2 | 0.016 |
| Gammaproteobacteria | 13 | f__Enterobacteriaceae | 0.52 | 0.0017 |
| | 46 | g__Aeromonas | 0.52 | 0.0017 |
| Gammaproteobacteria | 5 | f__Thiomicrospiraceae | 0.49 | 0.0017 |
| | 24 | o__Enterobacterales | 0.49 | 0.0017 |
| | 42 | f__Cellvibrionaceae | 0.28 | 0.0017 |
| Actinobacteriota | 8 | g__Micromonospora | 0.61 | 0.0017 |
| Cyanobacteria | 15 | f__Cyanobiaceae | 0.63 | 0.0017 |
| | 36 | f__Cyanobiaceae | 0.41 | 0.0017 |
| | 38 | c__Cyanobacteriia | 0.36 | 0.0017 |
| Campylobacterota | 16 | o__Campylobacterales | 0.15 | 0.026 |
| Desulfobacterota | 41 | None | 0.25 | 0.026 |
| Gammaproteobacteria | 27 | g__Alcanivorax | 0.25 | 0.026 |
| | 26 | o__Pseudomonadales | 0.1 | 0.026 |
| Others | 28 | f__Sphingomonadaceae | 0.12 | 0.026 |
| Gammaproteobacteria | 19 | g__Polynucleobacter | 0.29 | 0.0017 |
| | 1 | g__Thioglobus | 0.25 | 0.0017 |
| | 12 | g__BACL14 | 0.25 | 0.0017 |
| | 34 | None | 0.22 | 0.0017 |
| Others | 28 | f__Sphingomonadaceae | 0.2 | 0.0017 |
| Alphaproteobacteria | 2 | c__Alphaproteobacteria | 0.46 | 0.026 |
| Gammaproteobacteria | 19 | g__Polynucleobacter | 0.22 | 0.026 |
| Firmicutes | 45 | f__Clostridiaceae | 0.55 | 0.0059 |
| Alphaproteobacteria | 14 | g__Sinorhizobium | 0.35 | 0.0017 |
| | 30 | f__Rhodobacteraceae | 0.34 | 0.0017 |
| | 9 | f__Rhodobacteraceae | 0.23 | 0.0017 |
| | 40 | c__Alphaproteobacteria | 0.23 | 0.0017 |
| Thermotogota | 18 | g__Thermosipho | 0.3 | 0.0017 |
| Bacteroidota | 6 | c__Bacteroidia | 0.51 | 0.0017 |
| | 10 | c__Bacteroidia | 0.31 | 0.0017 |
| Deinococcota | 32 | f__Marinithermaceae | 0.34 | 0.016 |
| Thermotogota | 18 | g__Thermosipho | 0.34 | 0.016 |
| | 39 | g__Thermotoga | 0.2 | 0.016 |
| Bacteroidota | 10 | c__Bacteroidia | 0.33 | 0.028 |
| | 29 | None | 0.3 | 0.028 |
| | 6 | c__Bacteroidia | 0.15 | 0.028 |
| Firmicutes | 44 | o__Lactobacillales | 0.76 | 0.0017 |
| Gammaproteobacteria | 26 | o__Pseudomonadales | 0.41 | 0.0059 |
| | 3 | g__Fangia | 0.37 | 0.0059 |
| Spirochaetota | 4 | g__Spirochaeta_A | 0.57 | 0.035 |
| Gammaproteobacteria | 3 | g__Fangia | 0.81 | 0.0017 |
| Campylobacterota | 16 | o__Campylobacterales | 0.66 | 0.0032 |
| Gammaproteobacteria | 12 | g__BACL14 | 0.32 | 0.0032 |
| Thermotogota | 39 | g__Thermotoga | 0.66 | 0.0017 |

Supplementary Fig. 9: Strategies for B vitamins uptake associated to specific genome functional clusters (GFCs). ‘IndVal’ expresses the strength of the respective strategy-GFC associations and ‘p-adj’ is the false discovery rate adjusted *p*-value.

#### Broken vitamin pathways

Several genomes didn't have a complete biosynthetic pathway for at least one of the B vitamins and in a small portion of genomes the related transporter was missing too (~20%; Supplementary Fig. 10a). These problematic cases could reflect limitations of the annotation process (e.g. unknown transporters or alternative genes/pathways), however, there is growing evidence that organisms without complete biosynthetic pathways are able to grow on vitamin B intermediates [35,38]. Therefore, we looked for the presence of possible fragmented biosynthetic pathways that could suggest forms of auxotrophy towards specific B-vitamin intermediates. We found that nearly all genomes with problematic vitamin $B_1$ annotations possessed truncated biosynthetic pathways. However, while some of these genomes showed the capability to grow on exogenous precursors (e.g. pyrimidine moiety, HMP or the thiazole moiety, HET), others lacked the last enzyme of the pathway (Supplementary Fig. 10b). A few cases of genomes potentially relying on exogenous precursors were also identified for vitamin $B_7$ (e.g. d-desthibiotin) and vitamin $B_{12}$ (e.g. cobyrinic acid a,c-diamide or adenosylcobinamide). The rest of the problematic genomes possessed none or only a few annotated genes for such pathways (Supplementary Fig. 10c,d). These gaps could be due to limitations of the annotation step or to metabolic independence. Vitamin $B_1$ is a cofactor involved in several core metabolic processes (e.g. TCA cycle, amino acid metabolisms, pentose phosphate pathway) and the lack of the last enzyme may point to an annotation issue or to a specific adaptation of any $B_1$-dependent enzyme towards using the monophosphate version of vitamin $B_1$ (the last enzyme simply adds a second phosphate group). A similar conclusion could be drawn for vitamin $B_{12}$ and $B_7$, however there may be more support towards the metabolic independence. Vitamin $B_{12}$ is involved in amino acid and nucleotide synthesis, as well as in fatty- and amino acid breakdown, while vitamin $B_7$ is involved in a "side" path of the TCA cycle and in the urea cycle. Most of these processes have vitamin-independent routings (e.g. [39,40]) and some microorganisms are capable of $B_{12}$ independent growth [41] suggesting that in some cases these vitamins might not be essential.

s

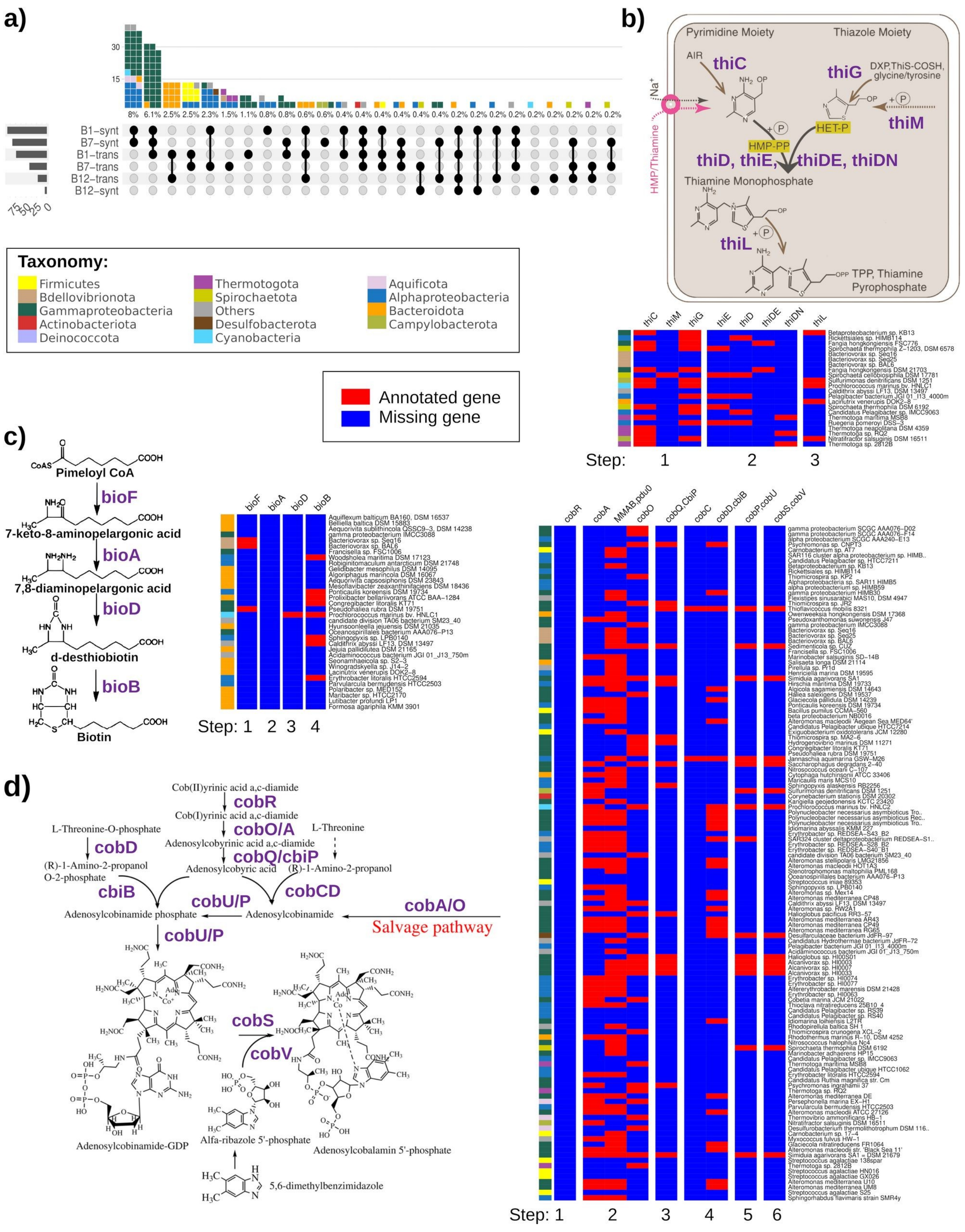


Supplementary Fig. 10: (a) Plots of intersecting sets showing incomplete combinations where both the biosynthetic pathway and the transporter for one (or more, 1% of genomes) of the B vitamin are missing; the left bar chart indicates the total number of genomes for each trait, the dark connected dots indicate the different configurations of traits and the waffle bar chart indicates the number (and percentage) of genomes provided with such a configuration; each piece of a waffle bar represents a genome and it is coloured according to the taxon. (b-d) Schematic overview of the biosynthetic pathways of (b) vitamin B1 (adapted from [44]; where HMP is the pyrimidine moiety and HET is the thiazole moiety), (c) B7 (adapted from [43]) and (d) B12 (adapted from [42]; the reaction for the biosynthesis of the tetrapyrrole compound are not shown). Presence-absence maps show the annotated and missing genes involved in the pathway for which a genome is missing both biosynthesis and transport capacities of a relevant vitamin.

### Siderophore and vibrioferrin traits' distribution

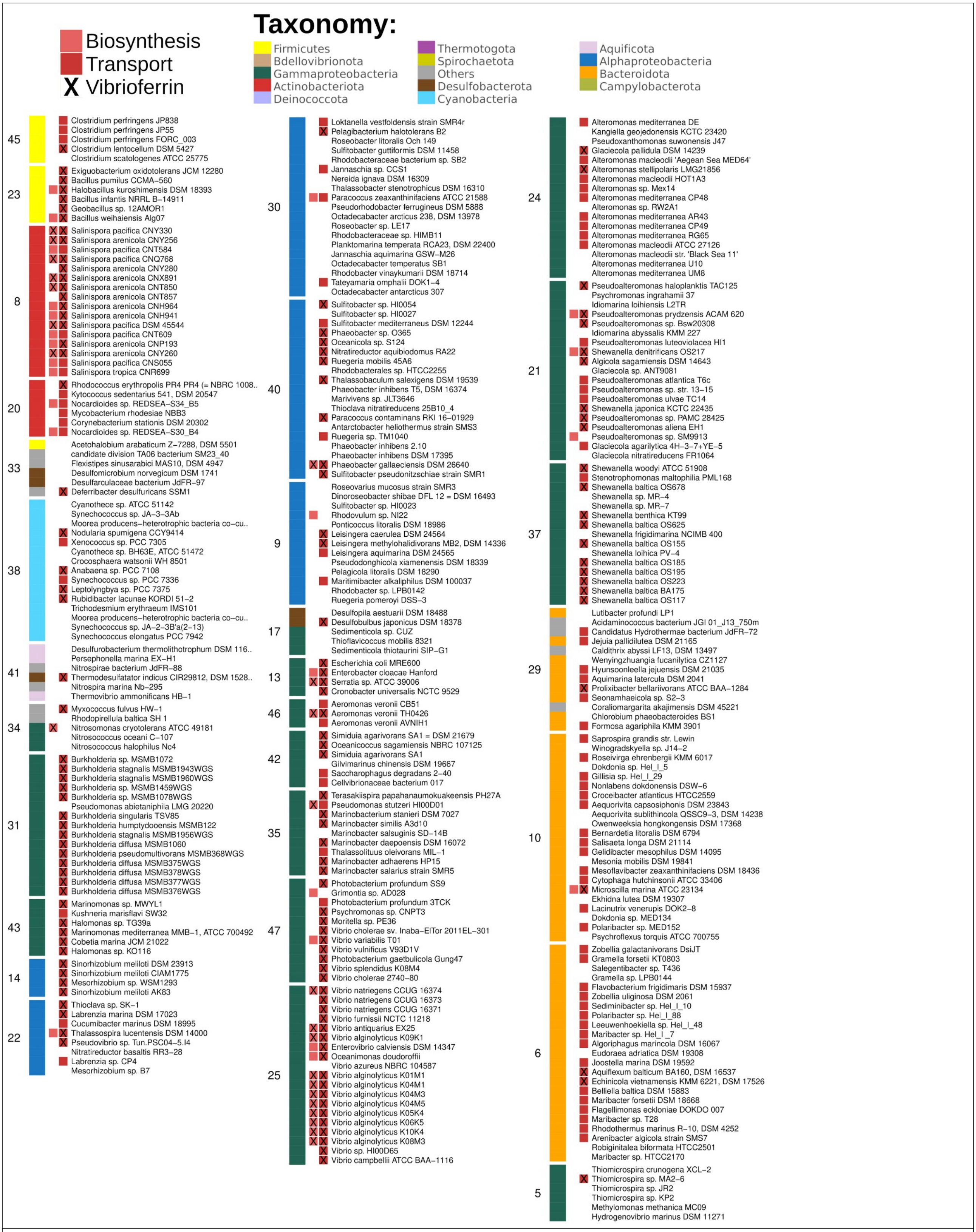


Supplementary Fig. 11: Distribution of siderophore biosynthesis and transport traits. Genomes are grouped in genome functional clusters (GFCs). Annotation of the specific vibrioferrin synthetic and transport operons is marked with an ‘X’.

#### Antibiosis traits’ distribution

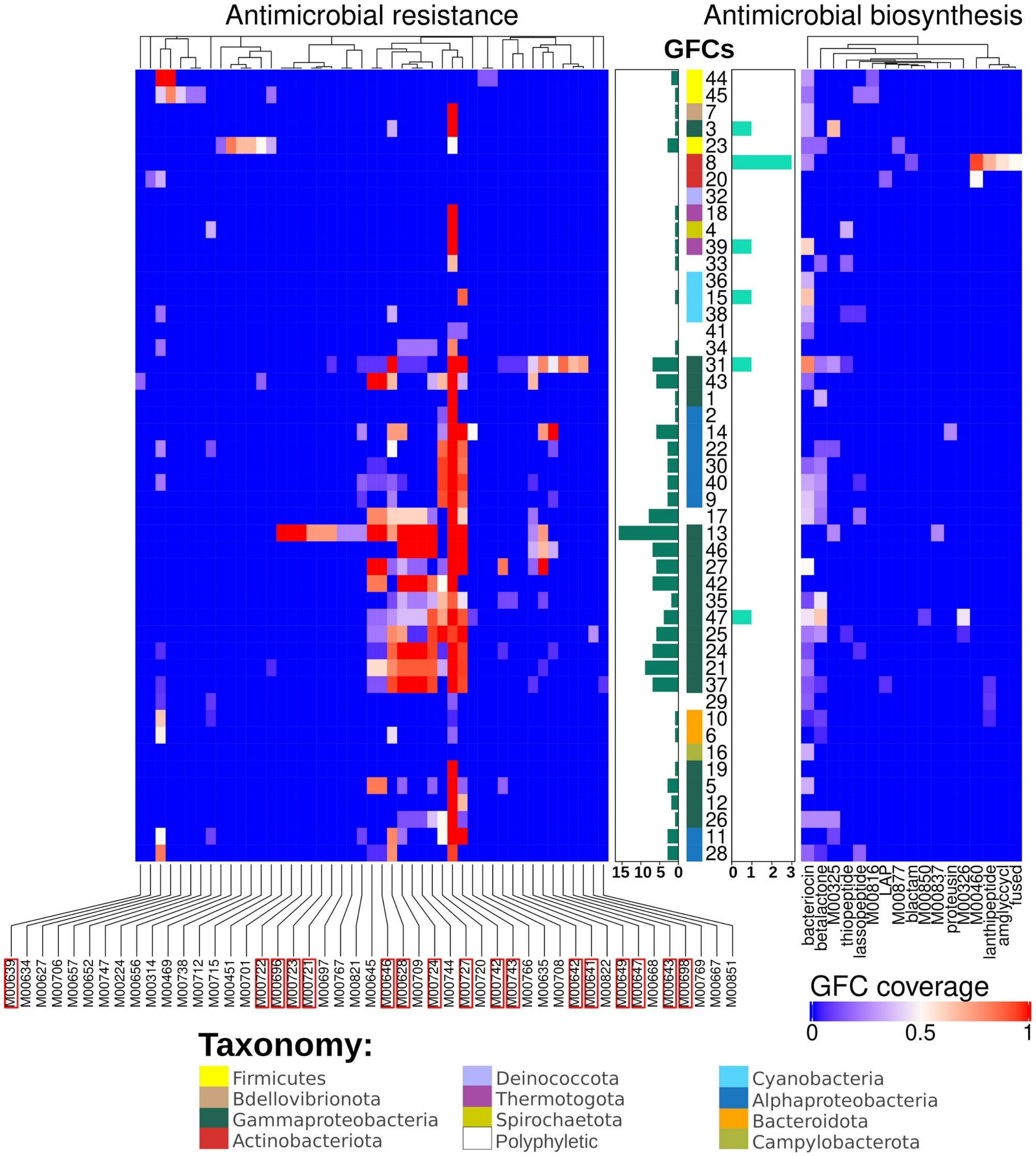


Supplementary Fig. 12: Distribution of antimicrobial resistance and biosynthesys traits in genome functional clusters (GFCs). Only traits that occurred in >50% of the genomes grouped in a GFC (i.e. GFC coverage) account for the trait richness in the central bar plots. Resistance traits marked with a red box are considered generic as the relevant KEGG modules are also involved in other cellular functions (e.g. cell division, protein quality control and transport of other compounds; see Supplementary Table 5 for details).

### Linked trait clusters (LTCs)

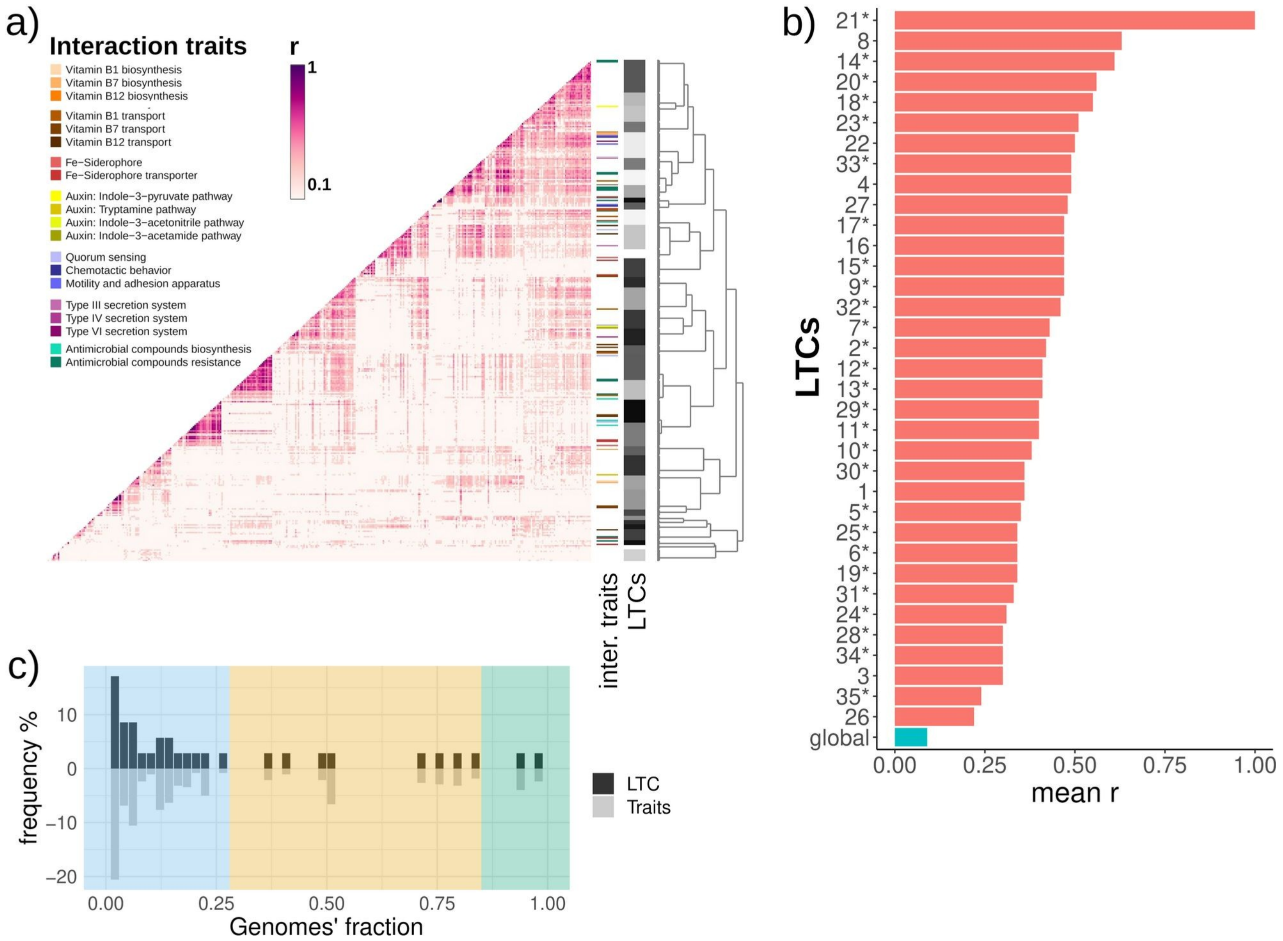


Supplementary Fig. 13: (a) r correlations among all genetic trait pairs; vertical colour bar indicates the presence of different interaction traits while the vertical grey bar delineates specific linked trait clusters (LTCs). An interactive version of the same figure is available as Supplementary file 2. (b) Mean r values of all pairs of genetic traits included in each LTC; mean r between all detected genetic traits (‘global’) is shown as indication of a random correlation within the dataset. (c) Histogram of LTC and genetic traits frequency across genomes’ fraction showing the division in “core” (present in ≥ 90% of genomes; green), “common”

(< 90% and ≥ 30%; yellow) and “ancillary” (≤ 30%; light-blue) LTCs.

#### Absence-pattern among common LTCs

Some of the genetic traits included in the common LTCs 2, 4 and 7 (found in 50-74% of the genomes) were consistently missing in some GFCs and/or taxonomic groups (Supplementary Fig. 14).

#### Broken TCA cycle

LTC 4 (mean r = 0.49) was absent in all Cyanobacteria (GFCs 15, 36 and 38), Thermotogota (GFCs 18 and 39), Spirochaetota (GFC 4) and most GFCs of Firmicutes (44 and 45)(Supplementary Fig. 14), which represented ~30% of genomes. It included three KEGG modules involved in the Citrate cycle: the complete TCA (M00009), the second carboxylation part (from 2-oxoglutarate to oxaloacetate; M00011) and the succinate dehydrogenase complex ($6^{th}$ reaction of the TCA cycle; M00149). These genomes, however, still had the first carboxylation part of this cycle (from oxaloacetate to 2-oxoglutarate; M00010) which was included in the core LTC 5.

For almost four decades it was thought that Cyanobacteria indeed was lacking a full TCA cycle, until two new enzymes were discovered. These enzymes catalysed together the conversion of 2-oxoglutarate to succinate and thus functionally replaced 2-oxoglutarate dehydrogenase and succinyl-CoA synthetase [45]. In our annotation, Cyanobacterial genomes lacked the two ‘classic’ reactions of the TCA cycle (M00009), and therefore the pathway was flagged as incomplete (based on our rule of one gap for traits with up to 10 reactions). A manual search revealed that the 2-OGDC gene (K01652), which catalyses the conversion of 2-oxoglutarate to succinic semialdehyde, was present in all Cyanobacterial genomes, whereas the SSADH gene (K00135), that catalyses the conversion of succinic semialdehyde to succinate, was present only in the genomes of Cyanobacteria belonging to GFC 38. All of the pico-Cyanobacteria genomes lacked the SSADH gene, consistent with the current view that these organisms lack a full TCA cycle [45].

In addition, our results showed that other heterotrophic bacteria lacked several or almost all of the reaction of this pathway. To the first case belonged the genomes of Spirochaetota and Thermotogota, (grouped in GFCs 4 and 39, respectively) while the latter case comprised several Firmicutes (GFCs 44 and 45) and the other Thermotogota (GFC 18) genomes. These findings broaden what was reported in previous studies which were focused on single species belonging to the mentioned taxa [46–49].

#### Differences in cell wall composition

The LTCs 2 and 7 (mean r = 0.42 and 0.43, respectively) were absent in 36-50% of the genomes and provided another direct way to compare and validate our findings as they included pathways related to the cell wall assembling. LTC 7 grouped genetic traits for the production of keto-deoxyoctulosonate (KDO; M00060, M00063 and M00866), a core constituent of lipopolysaccharide, and the lipopolysaccharide transporter (M00320; Lpt machinery). LTC 2 included genetic traits for the gamma−Hexachlorocyclohexane (M00669; similar to the *mla* machinery) and phospholipids (M00670; *mla* machinery) transport systems. All these traits, as well as transporters for lipoprotein (M00255; Lol machinery) included in LTC 4 described above, were consistently missing in GFCs 23, 44, 45 (Firmicutes), 8, 20 (Actinobacteriota), and 32 (Deinococcota; Supplementary Fig. 14). These absence-patterns could be largely explained by the bacterial cell wall type, as all these GFCs grouped gram positive bacteria which are completely missing the outer membrane. Therefore, they don’t require any KDO biosynthesis nor export of any lipoprotein or lipopolysaccharide out of the cell membrane [50]. They also don’t need to exchange any phospholipids between two membranes (with the *mla* machinery) [51]. Despite being gram-negative, Thermotogota (GFCs 18 and 39), Spirochaetota (GFC 4) and the pico-Cyanobacteria (GFCs 15 and 36) also exhibited similar absence-patterns. Experimental evidence supports the lack of lipopolysaccharide in Thermotogota [52] and of KDO in the cell wall of Cyanobacteria [53] and Spirochaetota [54]. However, the lack of any lipoprotein transporter in Cyanobacteria likely reflects an issue of the annotation, as these organisms are known to possess lipoproteins in the outer cell membrane [53].

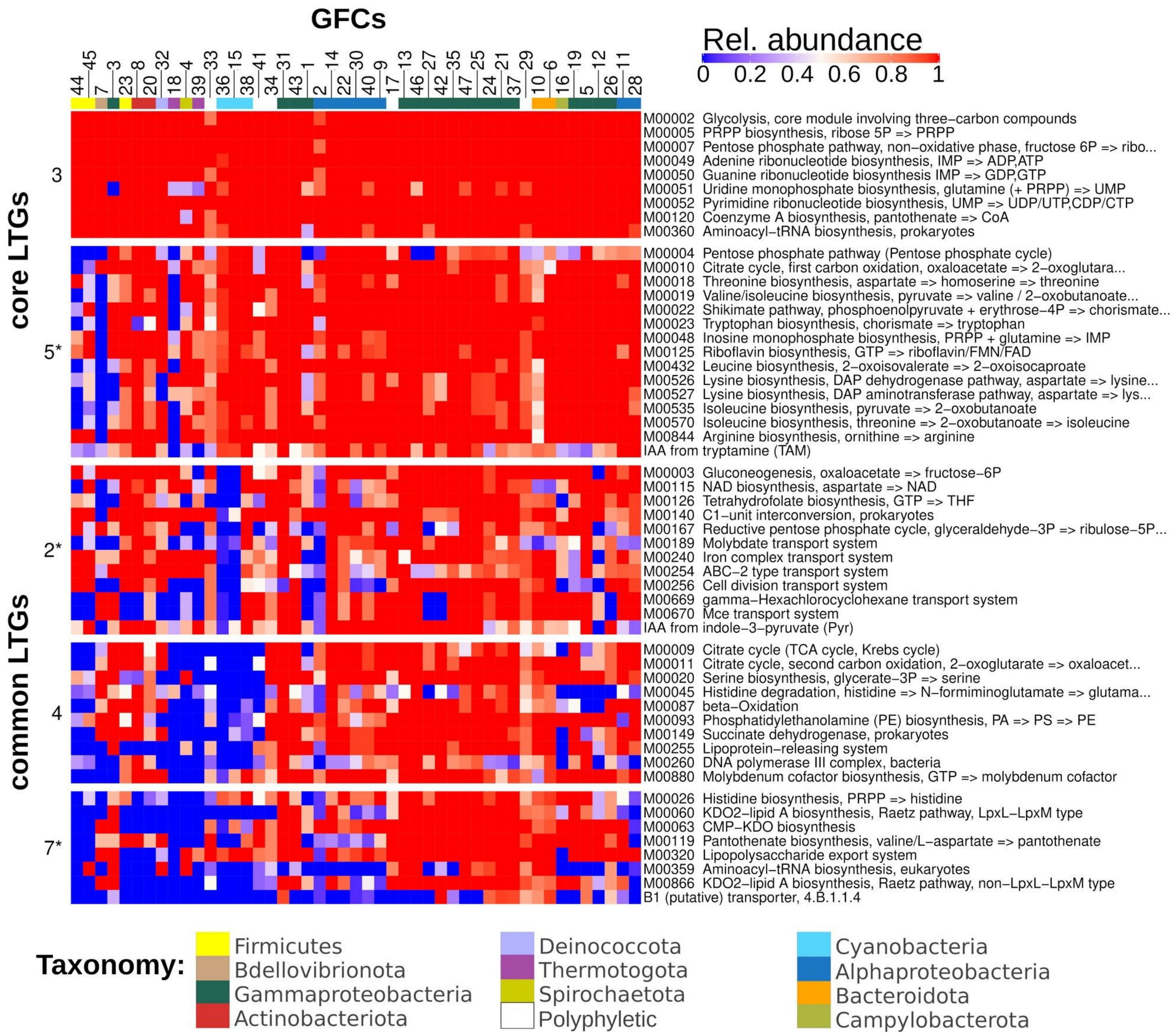


Supplementary Fig. 14: Relative abundance and absence-patterns of genetic traits grouped into core linked trait clusters (LTCs) 3 and 5, and in the common LTGs 2 (cell wall insight ), 4 (TCA cycle insight ) and 7 (cell wall insight) across all genome functional clusters (GFCs). The asterisks in LTC' names indicate that they also group an interaction traits. The horizontal color bar indicate the represented taxon in each GFC.

### Pipeline benchmarking

#### Criteria for KEGG module reconstruction

To assess the completeness of KEGG modules, we wanted to account for possible annotation issues (e.g. missing KOs due to the absence of suitable reference genes in the database, unknown genes, missannotations), therefore, we compared the outputs of different KM reconstruction analyses carried out using different thresholds (Supplementary Table 10). As there were no clear differences in the overall frequencies of complete KMs between the different tests (Supplementary Fig. 1c), we empirically determined the best criterion based on the expected absence/incompleteness of rare traits and the expected presence/completeness of core traits. For example, the most permissive threshold inspected (1 gap in KMs ≥ 2 reactions) allowed a KM of 2 reactions to be regarded as complete when only 1 reaction was annotated. While this criterion is arguably too permissive, it was discarded because it also predicted high frequency (0.38 – 0.71) of KMs for nitrate assimilation (anaerobic respiration in selected groups of heterotrophic bacteria [55]) and archaeal pentose phosphate pathway, which are expected to be nearly absent in an assembled genome dataset of pelagic marine bacteria. The next threshold (1 gap in KMs ≥ 3 reactions) was instead the best compromise, as it predicted a high frequency (0.43 – 0.95) of KMs mediating central processes such as the biosynthesis of uridine monophosphate and isoleucine, and other potentially relevant metabolisms like tetrahydrobiopterin biosynthesis (https://www.theseed.org/SubsystemStories/Pterin_biosynthesis/story.pdf) and glycogen degradation [17]. All these pathways are part of essential (e.g. synthesis of RNA, cofactors of central enzymes) and common cellular metabolisms that would never be found complete when applying the next threshold (1 gap in KMs ≥ 4 reactions). Moreover, out of the 491 complete KMs, only 167 showed changes in the frequency when compared with more stringent thresholds, but such differences were quite limited in magnitude (mean and median of the coefficient of variation were 42% and 21%, respectively).

#### Sensitivity analysis of clustering parameters

The advantage of clustering using the affinity propagation is that the algorithm automatically determines the best number of final clusters [56] without the need for the user to “guess” it a priori. This task is achieved by iterating through multiple clustering generated starting from different sets of initial exemplars, and by aiming to maximize the total similarity within each cluster. In the *apcluster* function (package apcluster; [57]), the parameter ‘q’ controls how the algorithm picks the exemplar nodes and affects the sensitivity of cluster detection. We therefore inspected how different q-values (ranging from 0 to 1) affected the results of the affinity propagation clustering for both GFCs and LTCs.

The cluster solutions were almost identical within the q-range 0.15-0.7 (for both GFC and LTC; Supplementary figure 15a-b), underpinning the robustness of this approach and of the detected clusters. We used q = 0.5 to run the final clustering for both GFCs and LTCs, as it was almost in the middle of such interval and it is the recommended values in the r package.

#### GFC & LTC clustering robustness

Using only high-quality and closed genomes, available (mainly) from cultured bacteria, inherently led to a skewed representation of certain taxonomic groups (see caption of Fig. 1). Therefore we tested the robustness of the detected GFCs and LTCs by down-sampling the most represented taxa, i.e. Gammaproteobacteria (34% of genomes), Alphaproteobacteria (22%), Cyanobacteria (15%) and Bacteroidota (11%). For each taxon, we randomly sampled 80%, 60% and 40% of the genomes 100 times and checked how often the genomes or the genetic traits were grouped in the same GFCs and LTCs, respectively.

Most GFCs always clustered together throughout each of the 100 starts and at all levels of down-sampling, while only 9 GFCs were found to be merged one with another GFC in some (20-50%) of the random starts (Supplementary figure 15c). Nevertheless, these events of merged-cluster did not specifically involved low abundant taxa (e.g. GFC 9 which included 13 genomes, GFC 40 with 19 genomes) and they mainly occurred in the taxa targetted from the down-sampling (the only exceptions were GFCs 34 and 41).

Similarly, also most of the LTCs clustered together throughout each of the 100 starts and at all levels of down-sampling. In some of the random starts, a few LTCs were found to be further split in smaller LTCs (i.e. LTCs 9, 11 and 17) or merged with another LTC (i.e. LTCs 18 and 30), however, these cases would not drastically change the traits' linkage of the original LTCs. In some of the random starts, a few genetic traits (~2-3) grouped in specific LTCs (i.e. 2, 4, 5, 6, 7, 19, 23, 28, 34 and 35) were found to be clustered in different

LTCs, suggesting that for these traits the functional linkage may be not consistently represented in the analysed genomes.

Overall, these results suggest that the over-representation of certain taxonomic groups did not specifically affect the clustering of GFCs and LTCs, underpinning the accuracy and reproducibility of the detected functional patterns.

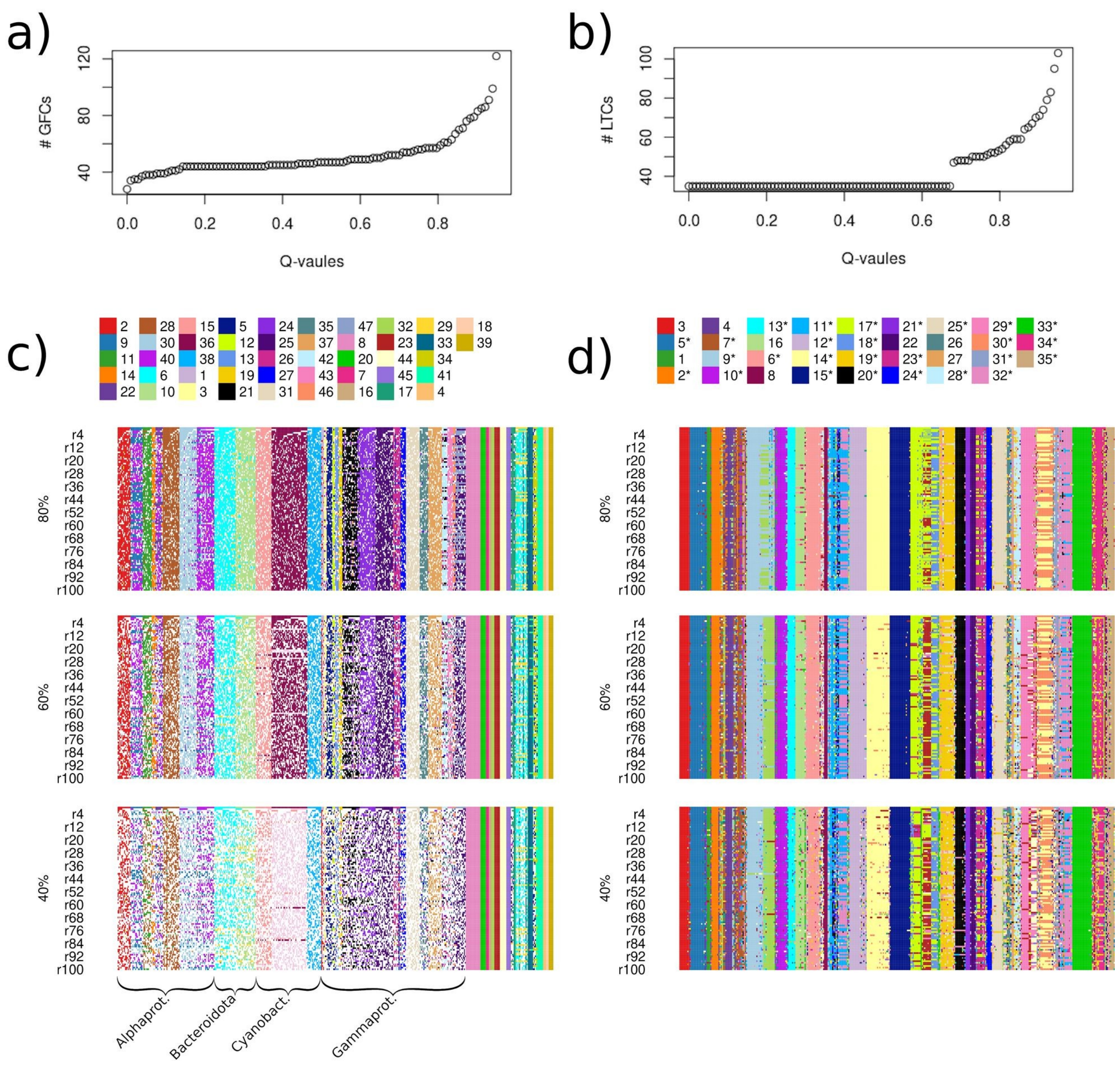


Supplementary Fig. 15: Sensitivity analysis showing the number of genome functional clusters (GFCs; a) and linked trait clusters (LTC; b) detected for different values of the 'q' parameter implemented with the *apcluster* function. Accuracy of GFCs (c) and LTCs (d) clustering assessed by performing 100 random down-sampling of the most represented taxa (i.e. Gammaproteobacteria, Alphaproteobacteria, Cyanobacteria and Bacteroidota) at 80%, 60% and 40% of their total genome counts. Down-samplings are shown as rows, while columns represent genomes (c) or genetic traits (d), and the different colours indicate the GFC or LTC membership.
